# Simulating the Influence of Conjugative Plasmids Kinetic Values on the Multilevel Dynamics of Antimicrobial Resistance in a Membrane Computing Model

**DOI:** 10.1101/2020.03.27.012955

**Authors:** Marcelino Campos, Álvaro San Millán, José M. Sempere, Val F. Lanza, Teresa M. Coque, Carlos Llorens, Fernando Baquero

## Abstract

Plasmids harboring antibiotic resistance genes differ in their kinetic values as plasmid conjugation rate, segregation rate by incompatibility with related plasmids, rate of stochastic loss during replication, cost reducing the host-cell fitness, and frequency of compensatory mutations to reduce plasmid cost, depending on the cell mutation frequency. How variation in these values influence the success of a plasmid and their resistance genes in complex ecosystems, as the microbiota? Genes are located in plasmids, plasmids in cells, cells in populations. These populations are embedded in ensembles of species in different human hosts, are able to exchange between them bacterial ensembles during cross-infection and are located in the hospital or the community setting, under various levels of antibiotic exposure. Simulations using new membrane computing methods help predict the influence of plasmid kinetic values on such multilevel complex system. In our simulation, conjugation frequency needed to be at least 10^−3^ to clearly influence the dominance of a strain with a resistant plasmid. Host strains able to stably maintain two copies of similar plasmids harboring different resistances, coexistence of these resistances can occur in the population. Plasmid loss rates of 10^−4^ or 10^−5^ or plasmid fitness costs ≥0.06 favor the plasmids located in the most abundant species. The beneficial effect of compensatory mutations for plasmid fitness cost is proportional to this cost, only at high mutation frequencies (10^−3^-10^−5^). Membrane computing helps set a multilevel landscape to study the effect of changes in plasmid kinetic values on the success of resistant organisms in complex ecosystems.

## Introduction

Plasmid kinetics are widely assumed to necessarily influence the spread of antibiotic resistance genes in bacterial populations and ecosystems (1-10). The main parameters that affect plasmid kinetics are: a) the rate of plasmid conjugation/transfer (the rate at which a bacterial cell harboring a conjugative plasmid [donor] transfers this plasmid to a recipient cell; b) the segregation rate due to plasmid incompatibility (considering the number of plasmid genome copies that are stably maintained in a bacterial cell); c) the rate of plasmid cost (the reduction imposed by the presence [and transfer] of a plasmid in the growth rate of the host bacterial cell); d) the rate of plasmid cost compensation (measuring the effect of mutations reducing plasmid cost); e) the frequency of mutational events in the plasmid or bacterial genome; and f) the rate of plasmid loss (the rate at which plasmids are lost during the bacterial replication process).

However, the effects of these changes on the kinetics of plasmid resistance genes among bacterial populations are necessarily influenced by numerous other factors acting in actual biological ecosystems, such as the intestinal microbiota. Of these factors, our previously published studies on modeling by membrane computing (11-13) considered the following: the ecosystem’s bacterial composition, the density and replication rate of cells in each species, their mutation frequencies, the content in chromosomal resistance genes, the selection intensity of resistant organisms due to differing antibiotic exposures, the elimination of susceptible bacterial populations by antibiotic treatment, the transmission of resistant bacteria among human hosts in hospital settings under differing admission-discharge rates, cross-colonization, and exposure to various antibiotics, as well as the influence of antibiotic resistance in non-hospitalized individuals who eventually pass through the hospital environment.

Experimentally investigating the influence of changes in plasmid kinetic parameters is extremely difficult and perhaps impossible, under real-world conditions. The research involves a complex, multilevel, multiparametric, and interactive landscape, involving genes, cells, populations, communities, hosts, and factors that influence transmission and selection. However, this problem can be approached using novel computational models integrating within-host and between-host modeling (14,15). Multilevel membrane computing models can provide an ecosystem-like framework composed of discrete independent but interactive units mimicking biological ones in a multi-hierarchical landscape of nested entities (e.g., genes inside plasmids, plasmids inside bacteria, bacteria inside microbiota, microbiota inside hosts, hosts inside the hospital, and interacting with the community) (12).

Membrane computing is conceptually based on complex biological systems, which are characterized by structured nested biological entities than can be conceptualized as separated by “membranes” (16,17). Each genes, plasmids, cells, species, populations, hosts, and compartments where the host is located (such as the community or hospital) is surrounded by “computational membranes” forming a multilevel nested structure, that can be studied by devices such as membrane systems or P systems (12, 13). The P system applied in this study mimics the complex biological landscapes in the computer world. In our model, each of the nested “membrane-surrounded entities” can independently replicate, propagate, become extinct, transfer into other membranes, exchange informative material according to flexible rules, mutate, and be selected by external agents (13). This computational model helps simulate the combined effect of changes in the various parameters, influencing the spread of antibiotic resistance plasmids. This study explores how changes in these parameters influence the spread of antibiotic resistance genes located in plasmids, in bacterial populations, and in microbial communities composed of different bacterial species, and which are the consequences of this spread. We also explore how these changes and their effects determine the frequency of various clones or species that harbor plasmids with antibiotic resistance genes. A simplified representation of the elements introduced in the membrane computing system is presented in Figure S1 (Supplemental Material).

In summary, we present a number of case studies analyzing the influence of changing plasmid kinetic values on complex hierarchical dynamics of antibiotic resistance in a simulated hospital setting. Most of the computing experiments referred to in this study mimics an evolution of 4.5 years (40,000 1-hour steps). To our knowledge, this is the first study that has addressed the effect of plasmid kinetic parameters in on the structure of multilevel biological systems.

## Results

The details and acronyms employed in the model are provided in Materials and Methods. However, to facilitate the understanding of the Results section, Table 1 is presented below.

**Table 1.**

| Acronym | Definition | Notes |
| --- | --- | --- |
| **AbA** | Antibiotic A | As aminopenicillins |
| **AbC** | Antibiotic C | As 3rd-generation cephalosporins |
| **AbF** | Antibiotic F | As fluoroquinolones |
| **AbAR** | Plasmid-mediated resistance to AbA | As TEM-1 |
| **AbCR** | Plasmid-mediated resistance to AbC | As ESBLs |
| **AbFR** | Chromosomal fluoroquinolone mutation | As *gyrA* mutation |
| **AbA\*R** | Chromosomal gene resistance to AbA in *K. pneumoniae* | As SHV-1 |
| **PL1** | Plasmid 1, originally in an *E. coli* population, with AbAR | Incompatible with PL3 |
| **PL3** | Plasmid 3, originally in *K. pneumoniae*, AbCR and AbAR | Incompatible with PL1 |
| **Ec0** | *E. coli* without plasmids, fully susceptible |  |
| **EcA** | *E. coli* with PL1 AbAR |  |
| **EcC** | *E. coli* with PL3 AbCR-AbAR |  |
| **EcF** | *E. coli* with chromosomal AbFR |  |
| **EcAC** | *E. coli* with PL1 and PL3 AbAR-AbCR |  |
| **EcAF** | *E. coli* with PL1 AbAR and chromosomal AbFR |  |
| **EcCF** | *E. coli* with PL3 AbCR-AbAR and chromosomal AbFR |  |
| **EcACF** | *E. coli* with PL1 AbAR, PL3 AbCR-AbCR, and AbFR |  |

### Influence of plasmid conjugation rates

How to estimate plasmid transfer rates is not a trivial question (18). In this simulation, conjugation rates are expressed as the proportion of donor-recipient contacts that result in a random and reciprocal cell-to-cell *E. coli*-*K. pneumoniae* plasmid transfer in a given period. For instance, 10^−6^ per hour, indicates that 1 in 100,000 contacts or 1 million donor-recipient contacts resulted in random and reciprocal cell-to-cell *E. coli-K. pneumoniae* plasmid transfers per hour. Conjugation rates widely differ in different plasmid-host combinations (19-22) and also is heavily influenced by the abundance and well-mixing of interacting bacterial populations (23). Our model included 3 conjugation rates (10^−3^, 10^−6^, and 10^−9^) applicable to plasmids PL1 and PL3, harbored by either *E. coli* or *K. pneumoniae*. The rate of spontaneous plasmid loss (segregation) was 10^−5^, and the mutation frequency for plasmid cost compensatory mutations was 10^−5^ (mutants reduced to one half the cost of harboring plasmids). Other default values are as described in the Materials and Methods section. The results for the various *E. coli* phenotypes emerging from the plasmid transfer and mutational resistance to fluoroquinolones are presented in Figure 1. At first sight, a dramatic effect on the *E. coli* population structure is only observed at high plasmid transfer rates (10^−3^) (Fig. 1a, 1b). A first selective burst of AbAR (red line) is followed by the AbAR-AbFR phenotype (brown) (due to the mutational evolution to fluoroquinolone resistance of AbAR cells) and by the acquisition of PL1 (AbAR) by the AbFR cells. The acquisition of PL3 from *K. pneumoniae* occurs almost simultaneously (now, AbA*R-AbCR, light blue) and then from AbFR *E. coli* (now, AbAR-AbCR-AbA*R, AbFR, dark blue). Subsequently, the predominant populations are AbFR *E. coli* harboring PL3 (AbCR-AbA*R, AbFR) (green).

**Figure 1:**
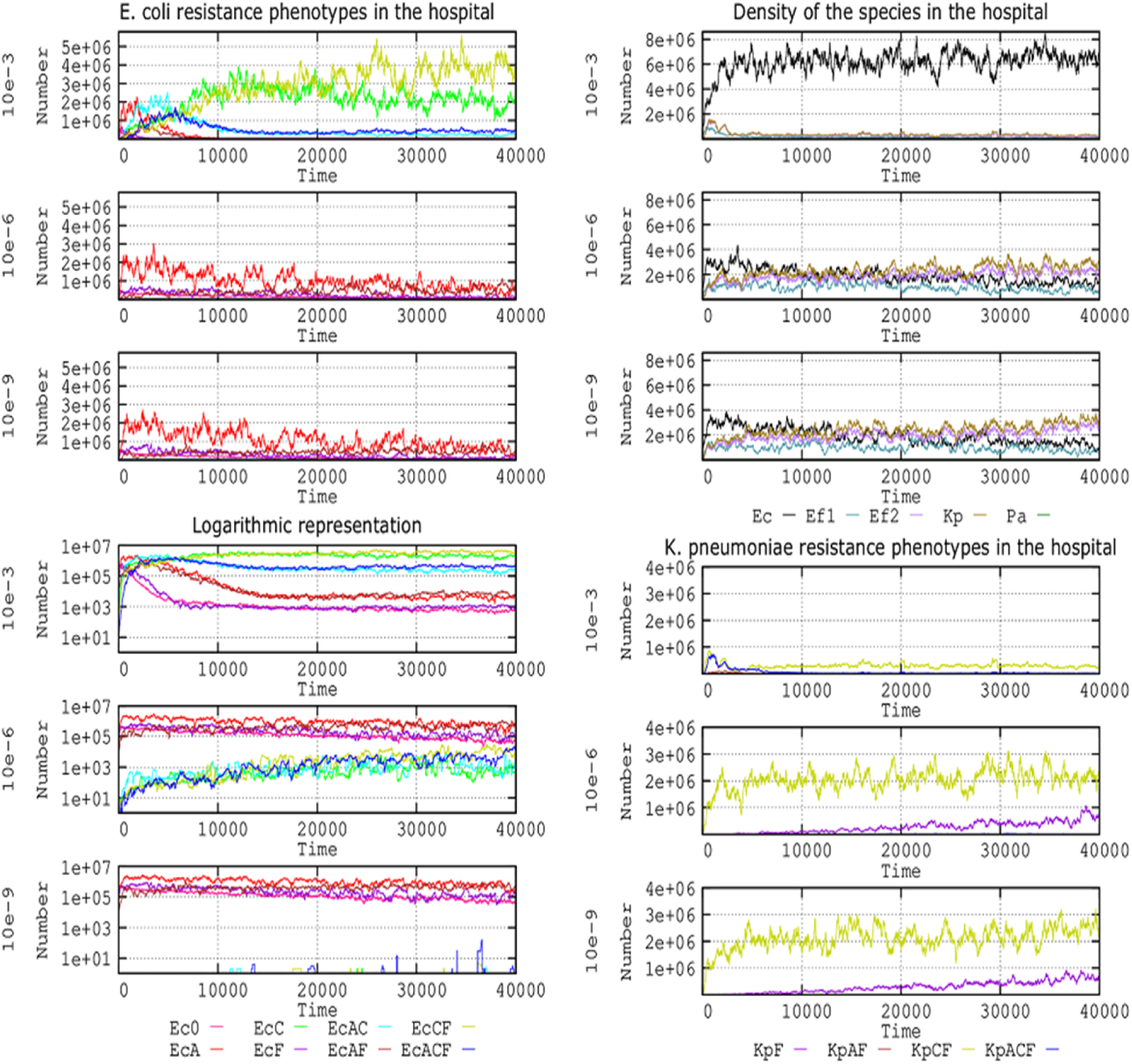
Influence of plasmid conjugation frequency (10^−3^, 10^−6^, 10^−9^) on the evolution of E. coli resistance phenotypes in the hospital. Upper left: Ec0, susceptible, no resistance plasmids (pink line), EcA, PL1-AbAR (red); EcC, PL3-AbAR-AbCR (light fluorescent green), EcF, AbFR (violet), EcAC, PL1-AbAR plus PL3-AbAR-AbCR, (light blue), EcAF, PL1-AbAR plus AbFR (brown), EcCF, PL3, AbAR-AbCR plus AbFR (olive green), EcACF, PL1-AbAR plus PL3-AbAR-AbCR plus AbFR (dark blue). Lower left: a logarithmic representation of the same resistance phenotypes. Upper right: density of the species E. coli (black line), K. pneumoniae (olive green), ampicillin-R E. faecium (violet), and ampicillin-S E. faecium (blue-green). Lower right: detail of the evolution of K. pneumoniae (olive green) and E. faecium. Numbers in ordinates are expressed in hecto-cells (one unit=100 cells in the microbiota)

*K. pneumoniae* is critical at the start of the process by providing the plasmid PL3 (AbCR-AbA*R) to *E. coli*; however, most plasmid transfers occur among the *E. coli* bacteria, thereby acquiring most of the antibiotic resistance benefits (Fig. 1c). With medium to low transfer rates, *E. coli* populations appear to remain stable and do not spread significantly in the human host population (lower boxes in Fig. 1a, 1c). Interestingly, a logarithmic representation (Fig. 1b) reveals remarkable differences at the 10^−6^ and 10^−9^ transfer rates in regard to the population structure. At both 10^−6^ and 10^−9^ transfer rates, there is a steady preservation of the fully susceptible *E. coli* population (pink), AbFR *E. coli* (violet) and containing PL1(AbAR) (red), and populations harboring PL1 (AbAR-AbFR) (brown). At the 10^−6^ transfer rate, however, a small part of the *E. coli* population acquires the PL3 plasmid from *K. pneumoniae* (and later from *E. coli*/PL3), giving rise to a constant increase in the phenotypes AbCR-AbA*R (green), AbCR-AbA*R-AbFR (olive green), and AbAR-AbCR-AbA*R, AbFR (dark blue).

Even at the 10^−9^ transfer rate, a few *E. coli* capture the PL3 plasmid but are unable to spread AbCR efficiently in the *E. coli* population. The long-term maintenance of *K. pneumoniae* with PL3 (olive green) is only assured at low transfer rates (10^−6^, 10^−9^), impairing the dominance of resistant *E. coli* populations (Fig. 1d), given that, at high transfer rates, the *E. coli* invade the *K. pneumoniae* cells with plasmid PL1, which might lose their resident PL3 plasmids due to incompatibility.

### Influence of plasmid incompatibility

Our model included the term “plasmid incompatibility” to explore the segregation of replicons sharing similar partitioning (par) loci, thus leading to mutual interference (24). We examined 2 highly related conjugative plasmids, PL1 and PL3. Conjugative plasmids typically have a limited number of plasmid copies (a maximum of 1 to 3 per cell, maximum 10) (25). In fact, there is an incompatibility phenomenon between replicons: the failure of 2 highly similar co-resident plasmids to be jointly inherited in a stable manner. This failure is due to the plasmids’ competition for replication factors or to delayed plasmid replication after the plasmid segregation in daughter cells (26,27). We examined the effect of cells tolerating only 1 plasmid copy (PL1 or PL3), where any new incoming plasmid copy (either PL1 or PL3) is rejected or substituted by one of the resident copies. We also examined the condition where only 2 plasmid copies can coexist (2 PL1s or 2 PL3s; or 2 PL1 and 2 PL3) and when 3 plasmid copies can coexist (all 3 PL1 or PL3; or 2 PL1 plus 1 PL3; or 2 PL3 plus 1 PL1). The question is particularly relevant, given that the presence of plasmid PL1 (AbAR) (originally located in *E. coli*) might prevent or influence the acquisition of plasmid PL3 (originally located in *K. pneumoniae*) or vice versa. **Figure 2** shows the results of the effects of these maximum plasmid copy numbers in the population structure of *E. coli* in the hospital setting.

**Figure 2.**
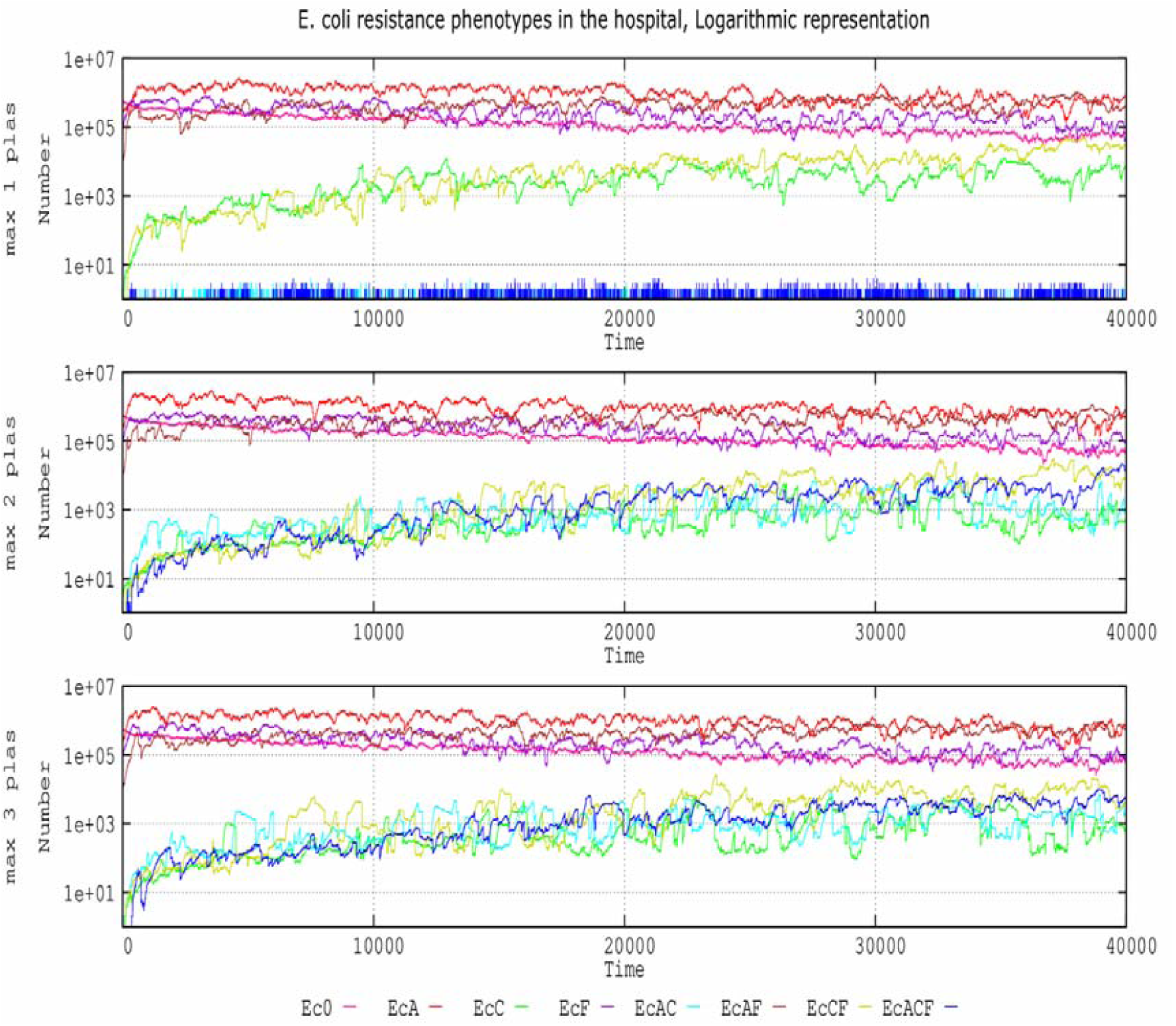
Effect of plasmid incompatibility on the evolution of E. coli antibiotic resistance phenotypes in the hospital setting. Ec0, susceptible, no resistance plasmids (pink line), EcA, PL-AbAR (red); EcC, PL3-AbAR-AbCR (light fluorescent green), EcF, AbFR (violet), EcAC, PL1-AbAR plus PL3-AbAR-AbCR, (light blue), EcAF, PL1-AbAR plus AbFR (brown), EcCF, PL3, AbAR-AbCR plus AbFR (olive green), EcACF, PL1-AbAR plus PL3-AbAR-AbCR plus AbFR (dark blue). The rate of plasmid cost compensation was fixed at 10-5. Numbers in ordinates are expressed in hecto-cells (one unit=100 cells in the microbiota)

When a single copy of the plasmid replicon is tolerated in the cell (originally the plasmid-bearing *E. coli* cells have a copy of PL1), there is a progressive invasion of E. coli cells by the plasmid PL3 from *K. pneumoniae*, giving rise to an increase in the cephalosporin-resistant (and ampicillin-resistant) phenotype EcC (green line). Through mutation of these cells, the phenotype also includes the fluoroquinolone-resistant phenotype (olive green), which can also arise with even higher frequency by the acquisition of PL1 by AbFR cells and then by displacement of PL1 by PL3 (initially in *K. pneumoniae*). These cells lose PL1 (AbAR); even if the PL1 plasmid is transferred, it does not remain in the cell (blue spikes at the bottom curve). With a maximum of 2 or 3 plasmid copies per cell, we can obtain a similar increase in resistant populations harboring both PL1 and PL3, with or without fluoroquinolone resistance (dark and light blue, respectively).

### Influence of random plasmid loss

In the process of cell division, the daughter cell receives at least one plasmid as a result of random diffusion influenced by multimer resolution systems (28). In our model, plasmid loss produced plasmid-free cells. The rate of spontaneous plasmid loss remains controversial (29). Our results (Fig. 3) indicate that at loss (random segregation) rates of 10^−3^, *E. coli* populations with plasmid PL1 or PL3 are not maintained beyond 8000 steps (approximately 2 weeks). At segregation rates of 10^−4^, the most abundant plasmids (PL1 in *E. coli*, with AbAR, in red) are maintained. After an initial increase, however, the plasmid population containing PL3 steadily decreases because of the incompatibility of PL3 with the dominant PL1 and due to the progressive reduction of *K. pneumoniae* with PL3, reducing the flow toward *E. coli*. Of course, cells with PL3 are more effectively selected than those with PL1, but PL1 is comparatively more abundant than PL3 in the ecosystem; therefore, PL1 is maintained along the 40,000 steps.

**Figure 3.**
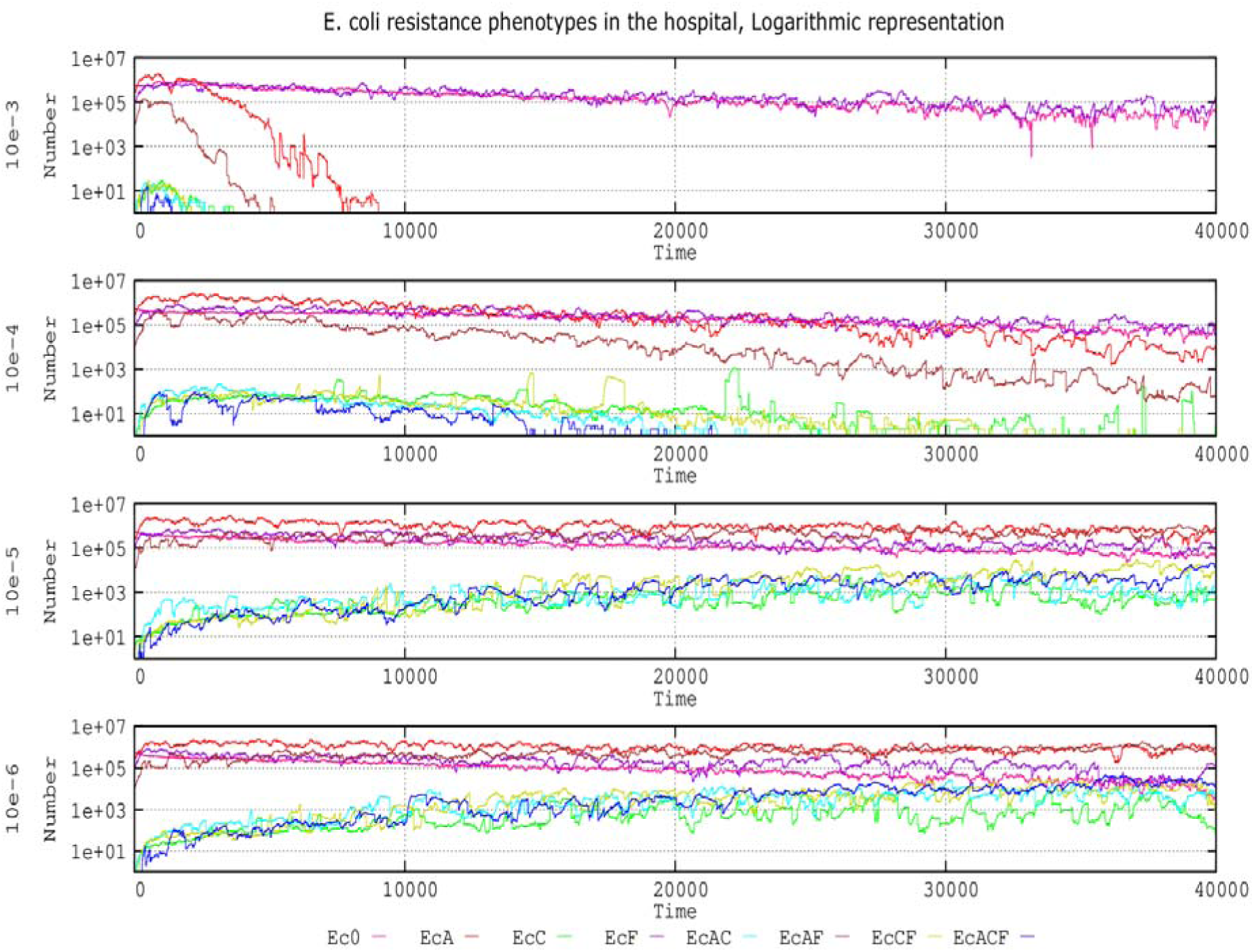
Influence of plasmid loss rates on the evolution of E. coli resistance phenotypes in the hospital environment. Ec0, susceptible, no resistance plasmids (pink line), EcA, PL1-AbAR (red); EcC, PL3-AbAR-AbCR (light fluorescent green), EcF, AbFR (violet), EcAC, PL1-AbAR plus PL3-AbAR-AbCR, (light blue), EcAF, PL1-AbAR plus AbFR (brown), EcCF, PL3, AbAR-AbCR plus AbFR (olive green), EcACF, PL1-AbAR plus PL3-AbAR-AbCR plus AbFR (dark blue). Compensation rate for plasmid fitness costs was fixed ad 10E-5. Numbers are expressed in hecto-cells (one unit=100 cells in the microbiota)

*E. coli* cells containing plasmid PL3 can slowly increase in number only at 10^−5^ segregation rates. At 10^−6^, these cells reach the density of antibiotic-susceptible or PL1 - containing cells. These results suggest that, at high plasmid segregation rates, the populations harboring the most abundant plasmids in the ecosystem have an advantage over populations with minority plasmids; however, if the loss of plasmids is relatively rare (such as 10^−6^), different plasmids might coexist in the population. Results with a 10^−^ 7 plasmid loss did not significantly differ from those of 10^−6^ (data not shown).

### Influence of plasmid fitness costs

Plasmid fitness costs imposed to the bacterial host are considered a factor that contribute for the spreading success of a particular plasmid and their genes (30-33). Several values for PL1 and PL3 plasmid fitness costs were included in the model to ascertain the effect of hosting these plasmids (or not) on antibiotic resistance and species composition. For reference, a plasmid fitness cost of 0.06 indicates a 6% reduced growth rate for the *E. coli* and *K. pneumoniae* strains harboring the PL1 or PL3 plasmid). We investigated the influence of these values with “no fitness cost” (fitness cost = 0). Default values included in this model were as follows: only 2 plasmids can coexist in a single cell; the rate of spontaneous plasmid loss is 10^−5^; the mutation rate to reduce 50% of the plasmid fitness cost is 10^−8^. The results are presented in Figure 4.

**Figure 4.**
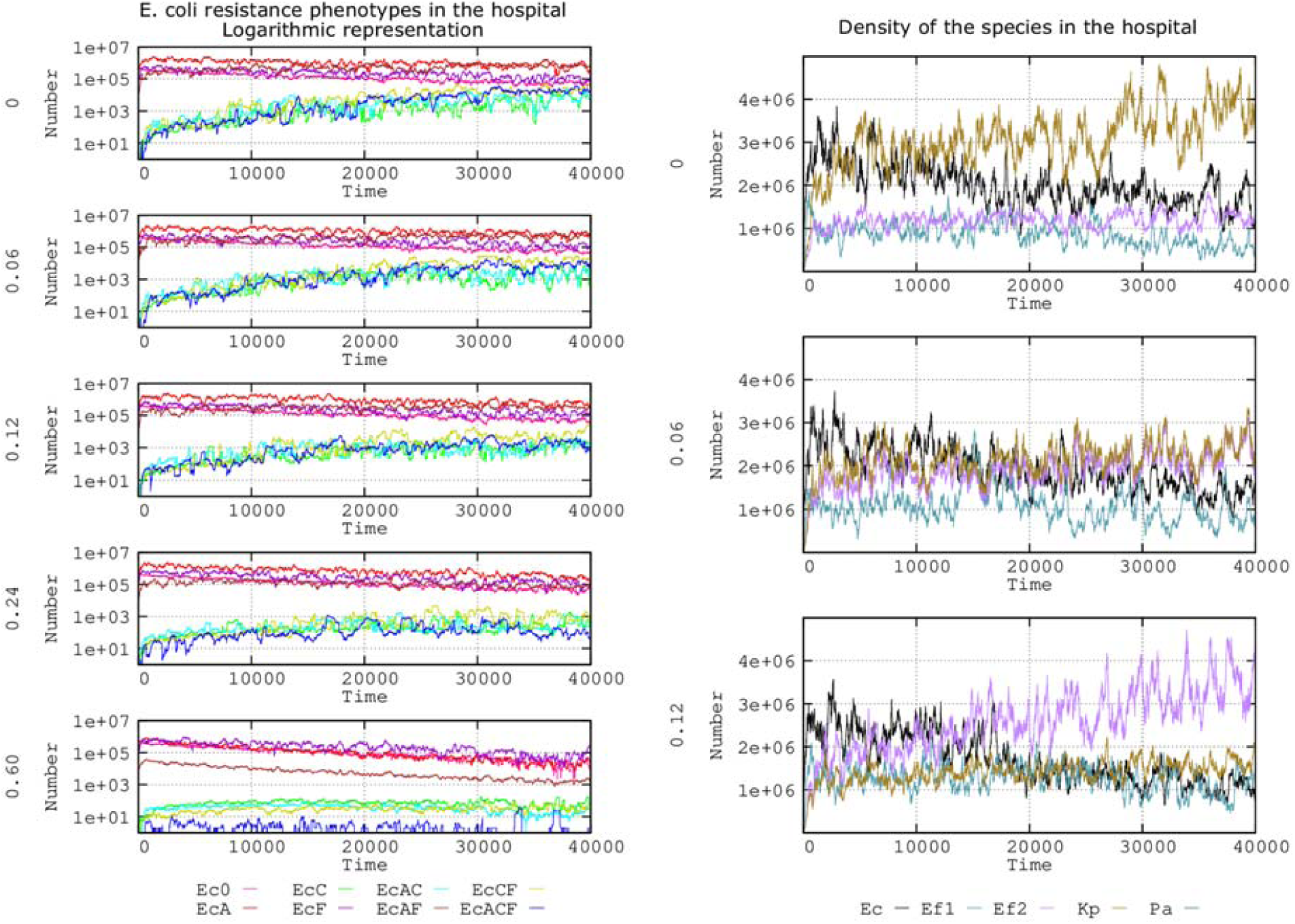
Influence of plasmid fitness cost on the evolution of E. coli antibiotic resistance phenotypes. The graphs in the left column show the effects for each fitness cost (0, 0.06, 0.12, 0.24, and 0.60). The lines indicate the following: Ec0, susceptible, no resistance plasmids (pink line), On the left part, effects of no fitness cost (0), and 0.06, 0.12, 0.24, and 0.6 fitness costs. Ec0, susceptible, no resistance plasmids (pink line), EcA, PL1-AbAR (red); EcC, PL3-AbAR-AbCR (light fluorescent green), EcF, AbFR (violet), EcAC, PL1-AbAR plus PL3-AbAR-AbCR, (light blue), EcAF, PL1-AbAR plus AbFR (brown), EcCF, PL3, AbAR-AbCR plus AbFR (olive green), EcACF, PL1-AbAR plus PL3-AbAR-AbCR plus AbFR (dark blue). The graphs in the right column show the influence of 3 fitness costs (0.0, 0.06, 0.12) on the species distribution in the simulated ecosystem: E. coli (black line), K. pneumoniae (olive green), ampicillin-R E. faecium (violet), ampicillin-S E. faecium (blue-green). Numbers in ordinates are expressed in hecto-cells (one unit=100 cells in the microbiota)

Regarding *E. coli* phenotypes (Fig. 4, left column) the effect of increasing the plasmid fitness cost was to steadily reduce the number of strains harboring only PL3 (encoding AbCR) or PL3 and PL1, alone or in combination with PL1 (AbAR) or AbFR (green, blue, olive green, dark blue lines). Only with a high plasmid fitness cost (0.60) was there a clear reduction of the predominant populations containing PL1 in the absence (red line) or presence (brown line) of chromosomal AbFR, probably due to the maintenance of an effective plasmid transfer. Note that the population with only chromosomal mutation (not influenced by plasmid fitness cost [AbFR] rises to dominance (violet line) but tends to decrease slightly, possibly because the reduction in cell multiplication provides reduced cell densities and therefore encourages the emergence of an AbFR mutation.

Differences in plasmid costs, even comparing no cost with 0.06 or 0.12 plasmid costs, might influence the species structure in the hospital ecosystem. At cost 0, *K. pneumoniae* with PL3 (olive green) steadily increases in frequency. *K. pneumoniae* originally benefits from the AbAR (including AbA*R) and AbCR phenotype and progressively acquires fluoroquinolone resistance, surpassing *E. coli* with PL1, with only the AbAR phenotype (black). At cost 0.06, the long-term dominance of *K. pneumoniae* is strongly reduced, but *E. coli* also decreases in frequency. With a cost of 0.12, *E. faecium* (in which PL1 or and PL3 are naturally absent but with a AbAR-AbCR phenotype [due to AbAR and AbCR chromosomal PBPs, violet]) tends to dominate. Thus, the cost of harboring a plasmid (decreasing growth rates) might influence the abundance and diversity of species present in a particular environments.

### Effect of changes in mutation frequency for compensation of plasmid fitness costs

Mutation frequency should influence the emergence of plasmid cost compensatory mutations. Compensatory mutations in the bacterial chromosome (30,34) and in the plasmid (35-38) might reduce the fitness cost of plasmid carriage, thus increasing the replication rate of the hosting microorganisms and the spread of plasmids. The emergence of compensatory mutations is certainly driven by the bacterial mutation frequency, but the effect of mutation might be asymmetrical if it occurs in the chromosome or plasmid, given that mutated plasmids can horizontally disseminate more effectively than mutated chromosomes through vertical transmission and given the presence of several genes that can compensate the cost, which differ among bacterial hosts (3, 36-43). In our basic model, high overall mutation were expected to influence not only plasmid cost compensation but also the selection of chromosomal mutants (for instance, fluoroquinolone-resistant mutants). To separate the two effects in our model and to detect the effect of compensating for plasmid fitness cost, we increased the mutation frequencies but maintained the basic mutation frequency (10^−8^) for AbFR observed for rifampicin (44) that can be applied for AbFR (45). In this model, fitness cost was established at 0.06, and the acquisition of a mutation reduces the fitness cost by a default value of 0.5. At a normal mutation frequency (10^−8^) or even 10^−5^ (data not shown), the effect of different strengths of mutational compensation on the frequency of plasmid-mediated antibiotic resistance was almost negligible in our model (**Fig. 5**).

**Figure 5.**
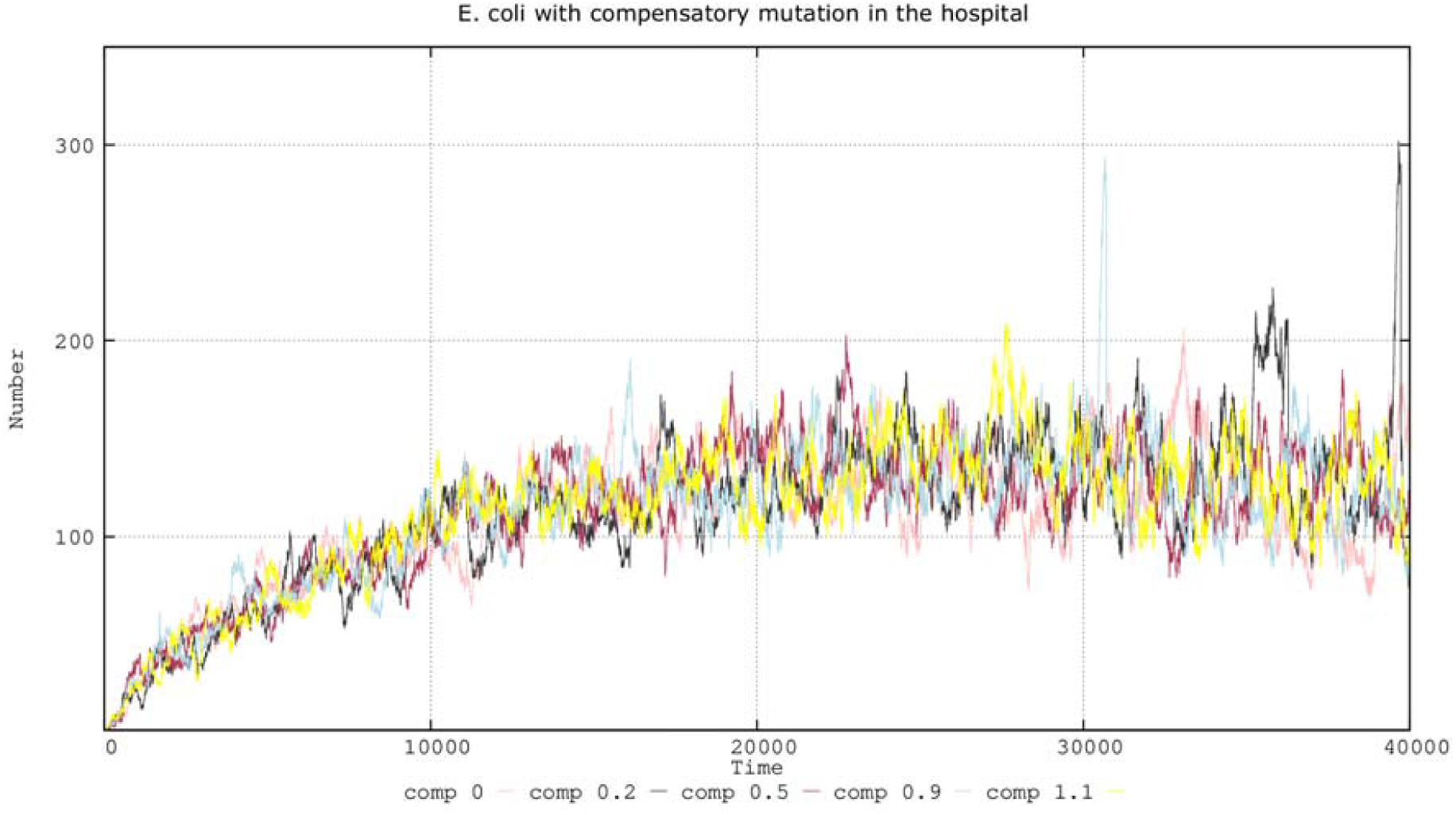
Effect of different compensation strengths of plasmid fitness cost. Lines: 0, no compensation, pink; 0.2 cost compensation, black; 0.5 cost compensation, brown; 0.9 cost compensation, blue; 1.1 cost compensation, yellow. Numbers in ordinates are expressed in hecto-cells (one unit=100 cells in the microbiota)

Figure 6 shows (left column) the effect of the 10-5 mutation frequency, which occurs by a small increase in *E. coli* lineages harboring compensated plasmids (olive green and dark blue lines), which is even more patent at the 10-3 mutation frequency. The reason for this small effect on *E. coli* can be explained by observing the overall landscape of the bacterial species included in the model (Fig. 6, right column); the proportion of compensated-plasmid-containing *K. pneumoniae* increases with the mutation frequency, probably at the expense of *E. coli* and *E. faecium*. Note that high mutation rates allow bacteria with initially low population sizes to cross the mutational threshold to obtain a beneficial mutation (46).

**Figure 6.**
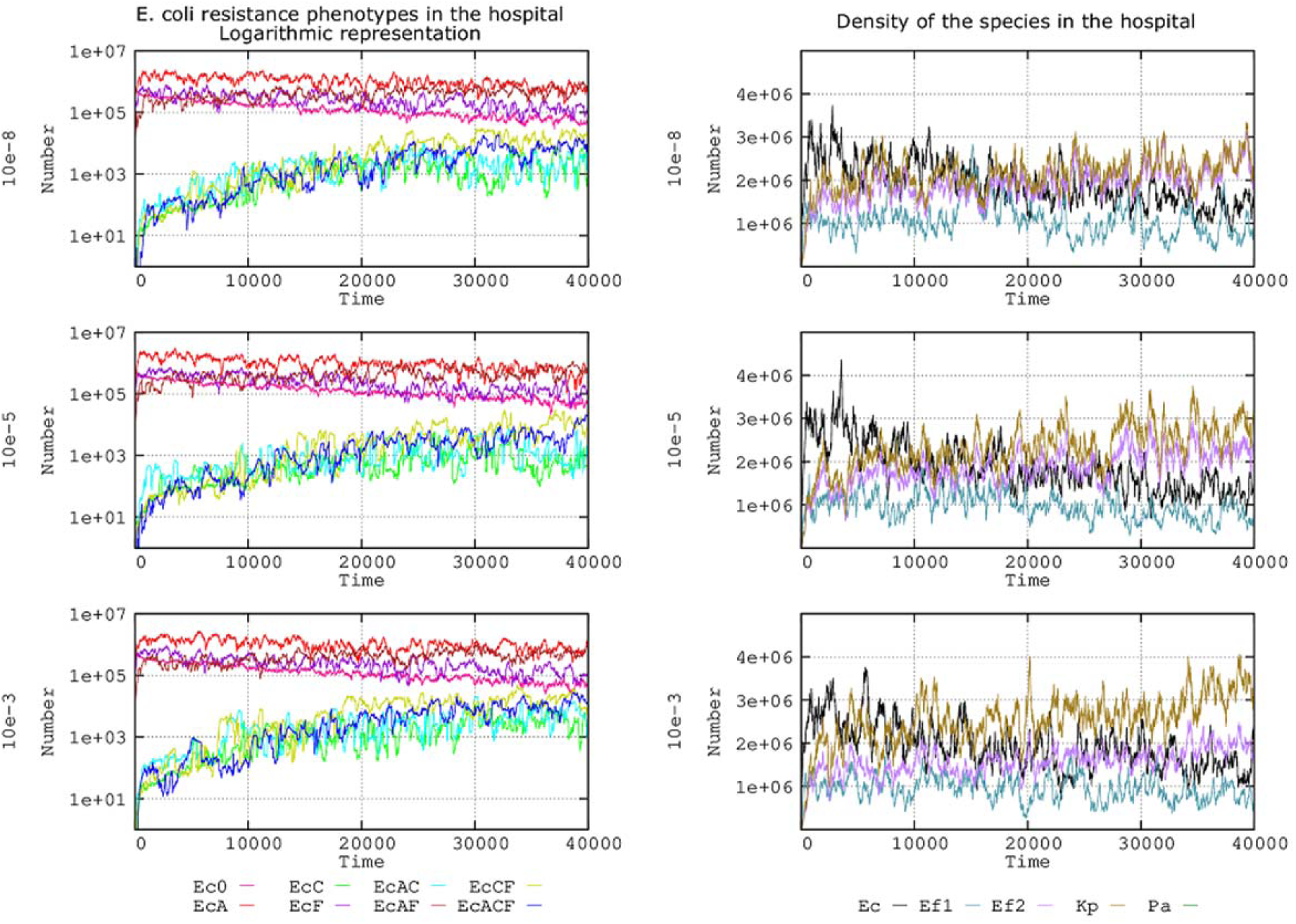
The left column shows the influence of different mutation frequencies (10^−8^, 10^−5^, and 10^−3^) compensating plasmid fitness costs in the evolution of E. coli resistance phenotypes. Ec0, susceptible to no resistance plasmids (pink line), EcA, PL1-AbAR (red); EcC, PL3-AbAR-AbCR (light fluorescent green), EcF, AbFR (violet), EcAC, PL1-AbAR plus PL3-AbAR-AbCR, (light blue), EcAF, PL1-AbAR plus AbFR (brown), EcCF, PL3, AbAR-AbCR plus AbFR (olive green), EcACF, PL1-AbAR plus PL3-AbAR-AbCR plus AbFR (dark blue). The right column shows the corresponding effect on the species composition, E. coli (black line), K. pneumoniae (olive green), E. faecium ampicillin-R (violet), ampicillin-S E. faecium (blue-green). Numbers in ordinates are expressed in hecto-cells (one unit=100 cells in the microbiota)

### Effect of changes in mutation frequency on combined fitness: compensation of plasmid fitness costs and acquisition of fluoroquinolone resistance

In the natural world, we can consider that different mutation rates might influence various bacterial functions differently. For instance, fluoroquinolone resistance mutations in topoisomerases typically occur at a rate of 10^−8^, but the frequency of mutations influencing reductions in plasmid fitness costs might be much higher (e.g., 10^−5^) (40). In the Figure 7 there is a representation of the evolution in our complex landscape of the number of cells with plasmid cost compensation with 10^−8^ or 10^−5^ fluoroquinolone resistance mutation frequency.

**Figure 7.**
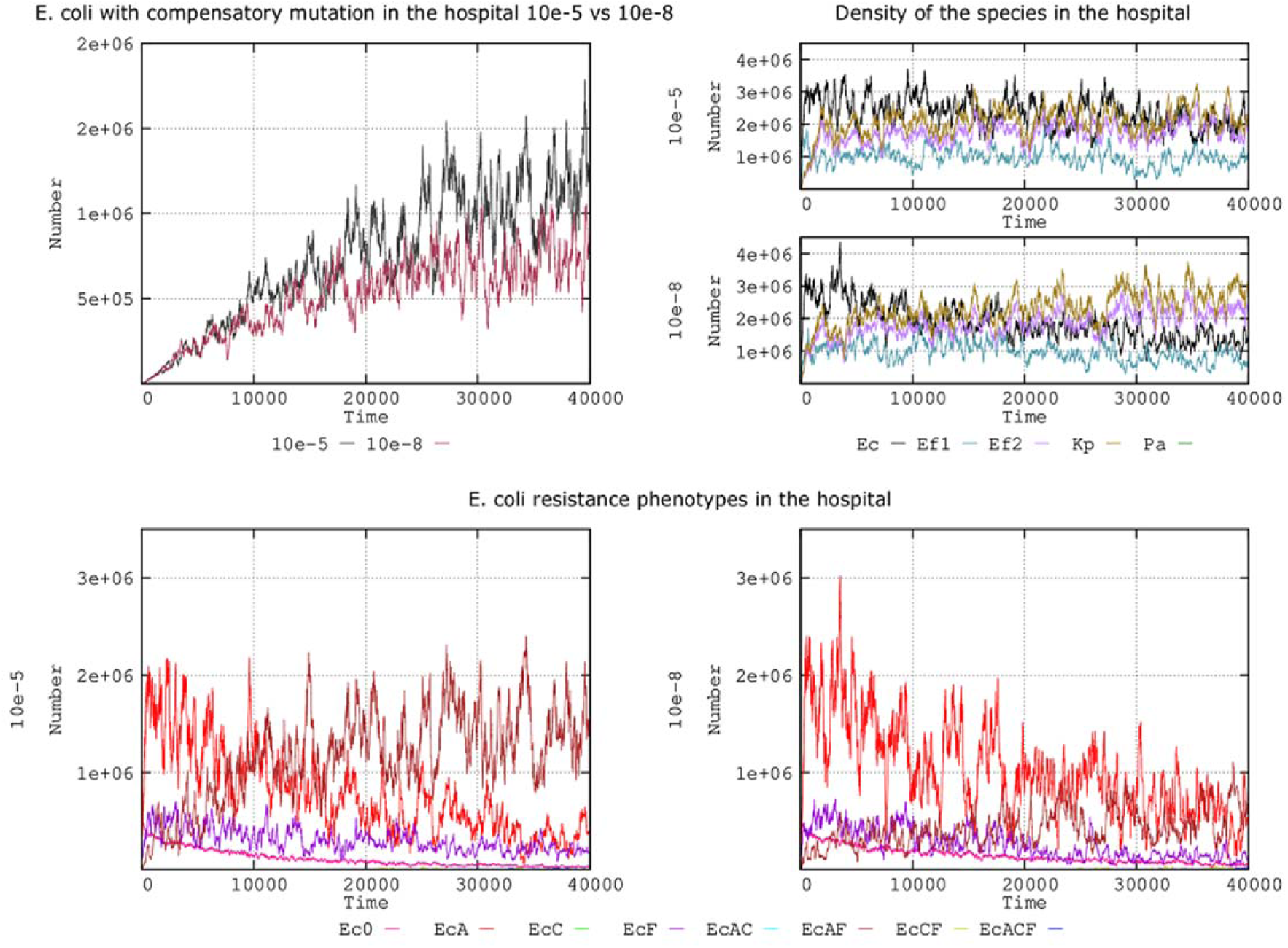
Compensation of plasmid fitness costs and frequency for fluoroquinolone mutation. Upper left: E. coli cells compensated for fitness costs when mutation rates for fluoroquinolone resistance are 10^−8^ (black) or 10^−5^ (brown-violet). Upper right: the effect of these mutation rates on species distribution (E. coli [black line], K. pneumoniae [olive green], ampicillin-R E. faecium [violet], ampicillin-S E. faecium [blue-green]. Bottom: evolution of E. coli antibiotic resistance phenotypes at fluoroquinolone mutation frequencies of 10^−5^ (left) and 10^−8^ (right). Numbers are expressed in hecto-cells (one unit=100 cells in the microbiota)

Statistically, cells with emerging mutations resulting in AbFR rarely compensate the plasmid costs; however, once an AbFR mutation selects an abundant resistant population (47), there is a higher probability that a mutation will emerge in this population and reduce the plasmid fitness cost. Figure 7 (up, left clumn) shows the evolution of the number of cells in our complex landscape with plasmid cost compensation when the mutation frequency for fluoroquinolone resistance was 10^−8^ or 10^−5^. This result indicates that (as stated above), *regardless* of the mutational frequency for plasmid cost compensation, the increased survival of fluoroquinolone-resistant cells in the hospital environment increases the absolute number of plasmid cost compensatory mutants. In the 2 panels of the first column of Figure 7b this effect is visible in the higher nosocomial prevalence of *E. coli* (black line) when AbFR emerges at a frequency of 10^−5^. This strong increase in the AbA-AbFR population carrying PL1 (AbA) in *E. coli* (brown line) when the mutation frequency is 10^−5^ is depicted in Figures 7 (left column). The increase in cost-compensatory mutations of PL1 in the increased AbA-AbFR population likely promotes its spread at these mutation frequencies, competing with the less frequent plasmid (PL3), resulting in no clear advantage for AbCR (data not shown, available on request).

### Combined effects of compensation values for plasmid fitness cost and variation in plasmid fitness cost

Figure 8, left column (experiment with fixed mutation frequency for plasmid fitness cost compensation of 10^−5^) shows that the number of plasmid cost-compensated *E. coli* is lower when the cost of carrying the plasmid increases (given that bacteria replicate more slowly); however, cost-compensated bacteria (green line) outpace the non-compensated bacteria (red line) much earlier. Thus, the expected benefit of mutational compensation is proportional to the fitness cost imposed by the plasmid (37). By increasing the plasmid fitness cost, the replication of bacteria is impaired, thus possibly reducing the population size and the possibility of obtaining compensatory mutations. Consequently, the reduction in plasmid fitness cost by compensatory mutations should increase population sizes. On the other hand, the benefit of high mutation frequencies in reducing the plasmid fitness cost should be proportional to this cost. Figure 8 (right column) shows the outcome for our “hospital scenario” of the *E. coli* population with a cost-compensated and a noncompensated plasmidome. For the same level of compensation (0.2), the rise in compensated cells (green line) is higher for 0.12 than for 0.06 plasmid fitness costs. If we increase the fitness cost compensation level (to 0.5), the rise in mutated cells is even higher, as expected, because the mutational yield is proportional to the population size.

**Figure 8.**
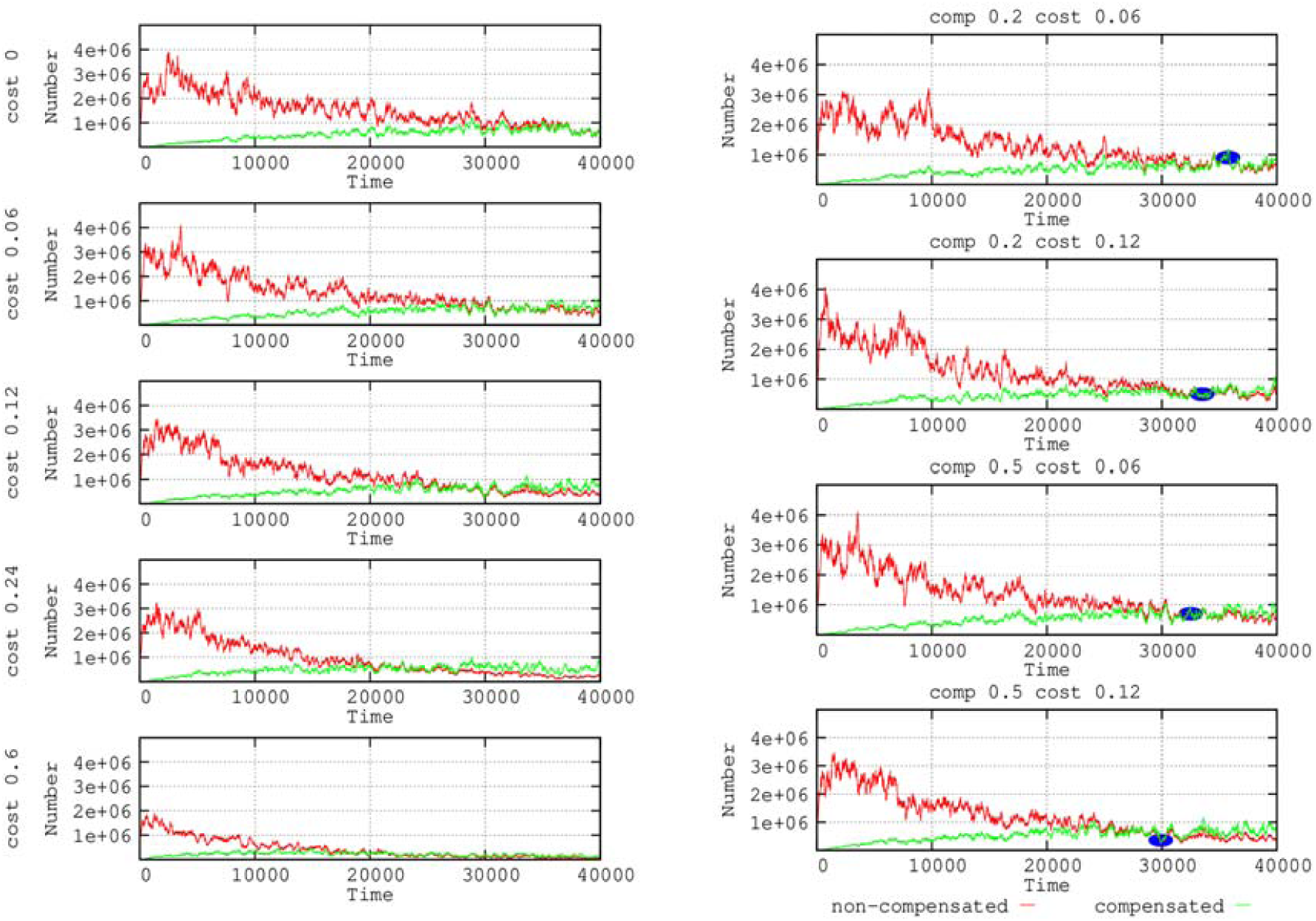
Effects of plasmid compensation depending on the plasmid fitness cost. In the left column, no compensated E. coli cells for plasmid fitness cost (green line) versus compensated cells (red). In the right column, combinations of 2 fitness cost values (0.06 and 0.12) with 2 strengths of compensation values (0.2 and 0.5). Small blue ovals highlight that when the plasmid cost is higher the compensation is more effective. Numbers in ordinates are expressed in hecto-cells (one unit=100 cells in the microbiota)

## Discussion

Plasmid biology should consider the multi-dimensional space where plasmids replicate and disseminate, not only inside and between bacterial cells, but in complex ecosystems. The application of membrane computing models to study the horizontal conjugative transfer of antibiotic resistance genes in bacteria (ref) is one of the few available approaches for addressing bacterial evolutionary dynamics in such a broad ecological context (14). Based on our previously published model (12), in this study the influence of various plasmid kinetic values in the evolution of antimicrobial resistance (8), was modeled within a complex system resembling the natural conditions that influence transmission at different levels (e.g., the flow of human hosts in the hospital and community, bacterial transmission/transfer rates among hosts, bacterial population sizes in the hosts, exposure and effects of various antibiotics in reducing bacterial numbers, selection of antibiotic resistant species, and the influence of “space for colonization” of resistant strains in the microbiota (12).

Details of the basic model’s design have been presented elsewhere (11, 12, 13). These integrative models are mostly fed with data on plasmid biology obtained through *in-vitro* experiments and suggest that predictions based only on laboratory data might not necessarily reflect the evolution of resistance in natural clinical landscapes. This study presents only a model under particular conditions (see Materials and Methods section), which were selected as representative examples; however, many other conditions can be introduced into the parameters in our accessible model (https://sourceforge.net/projects/ares-simulator/). Our main findings might help explain the relative weight of parameters that modify plasmid kinetics in the evolution of antibiotic resistance.

Plasmid transmission rates (the conjugation rates in our model) have been considered one of the main drivers of the spread of antibiotic resistance genes in natural bacterial populations (27). In fact, there is a line of research on ecology-evolution (eco-evo) drugs that seeks to develop plasmid conjugation inhibitors to reduce the burden of antibiotic resistance (48, 49). In our complex multilevel system and under the fixed conditions of the simulation, only high (10^−3^) conjugation rates clearly influence the dissemination of plasmids and the resistances they contain. However, we were able to differentiate between conjugation rates of 10^−6^ (in which the plasmid PL3 slowly propagates) from 10^−9^ (in which effects or transfers are no longer visible). Interestingly, higher conjugation rates (at which *E. coli* transfer PL1 [AbAR] effectively) tend to displace PL3 (AbCR, AbAR) resistance in *K. pneumoniae* (through *par* incompatibility). Thus, the PL3 is only preserved under low conjugation rate conditions in *K. pneumoniae.*

Plasmid incompatibility refers to the number of plasmids (replicons) sharing the same *par* system (in our case, PL1 and PL3) that can coexist in the same cell. In our simulation and under conditions allowing only a single plasmid to be maintained, PL3 (containing AbCR-AbAR) from *K. pneumoniae* can displace PL1 (AbAR); thus, AbCR significantly increases in *E. coli*. Conversely, the introduction of PL1 from *E. coli* into *K. pneumoniae* severely reduces the number of *K. pneumoniae* cells harboring PL3. In the conditions set in the present simulation (conjugation rate 10^−6^), if the cell tolerates 2 replicon copies and PL1 + PL3 can coexist, the number of *E. coli* cells harboring PL3 (AbCR) comparatively decreases, and there are no significant differences if the cell is able to maintain 3 replicons, a result that might be modified by the conjugation rate. In a previous published study by our group (12), *K. pneumoniae* retained PL3 (AbCR) if the conjugation rate is higher, 10E-4. Note that in the real world, *K. pneumoniae* frequently serves to introduce in the ecosystem plasmids with AbCR, that are then transferred to *E. coli*, and subsequently among *E. coli* populations (50, 51). Because *E. coli* has a comparatively higher population size, and most of the conjugations occur at the intraspecies level, in many cases the AbCR hospital “epidemics” tends to occur at long term in *E. coli*, and AbCR *K. pneumoniae* is usually maintained at a lower frequency (52,53,54,55).

Plasmids segregate (disappear, are lost) from the cells in which they are hosted, but the segregation rates are not well established and depend on the ecological-physiological conditions of the bacteria and the segregation mechanisms. In our model system, plasmid loss rates of 10^−3^ led to plasmid extinction. In general, high segregation rates favor plasmids contained in large bacterial populations. At 10^−4^, for example, only PL1, contained in the dominant *E. coli* population persists in time, displacing populations with PL3, despite the stronger selection pressure (due to AbCR). Higher exposure to antibiotics (cephalosporins) selecting for PL3 should logically favor the maintenance of PL3. In any case, PL3 persists much more effectively at low segregation levels (such as 10^−5^ and 10^−6^).

Plasmids fitness costs, expressed as a reduction in bacterial growth rate, has a relevant influence on bacterial resistance phenotypes, particularly on the propagation of plasmids hosted by minority populations. This effect is particularly visible when the plasmid cost is ≥0.12. For instance, the spread of PL3 (with AbCR) primarily hosted by *K. pneumoniae* is strongly reduced beyond a cost of 0.12. Even if the same cost was imposed by PL1, as is frequently present in the dominant *E. coli*, the reduction in fitness is somewhat compensated by intra-specific transfer; however, when the fitness costs is 0.60, the population with PL1 steadily decreases. Although AbFR populations are not influenced by plasmid fitness costs, many of the AbFR cells are lost because of the reduced reproductive rate imposed by the cost of plasmid they might contain.

Mutations might compensate the plasmid fitness costs. The mutation frequency should therefore affect the number of cost-compensated plasmids, particularly of the plasmids imposing a higher fitness cost. In our model, the effect on plasmid-mediated resistance was almost undetectable at the “consensus” mutation frequency (10^−8^), which might suggest that this compensation parameter has low epidemiological consequences. Even at a general mutation frequency of 10^−5^, the effects on the spread of antibiotic resistance under our experimental conditions were barely detectable. Of course, there are hypermutable strains (with a 10^−6^ or 10^−5^ mutation rate), and considering that the frequency of mutations influencing plasmid compensation might reach 10^−5^ (40), we cannot reject the possibility of scenarios in which the mutational cost compensation of *E. coli* lineages harboring compensated plasmids tends to increase (56). In fact, this might explain why *E. coli* isolates harboring plasmids with extended-spectrum β-lactamases have increased mutation frequencies (57). As proof of this concept, the benefit for cost-compensated strains (and for *K. pneumoniae*) becomes clear in a hypothetical scenario containing strains with a 10^−3^ mutation frequency (37).

Given that increases in general mutation frequency should necessarily influence the emergence of chromosomal fluoroquinolone-resistance mutations (AbFR), we combined the effects of the mutation rate on AbFR acquisition and plasmid-cost compensatory mutation. Our results indicate that, regardless of the mutational frequency for plasmid cost compensation, the increased survival and population increase of fluoroquinolone-resistant cells with high mutation rates increase the absolute number of plasmid-cost compensatory mutants.

In principle, the beneficial effect of plasmid cost compensation on the evolution of plasmid spread should be proportional to the reduction in fitness imposed by plasmid carriage. On one hand, our simulation shows that the cost-compensated bacteria surpass the number of non-compensated bacteria earlier when the cost is high. However, the number of plasmid cost-compensated *E. coli* decreases when the cost of carrying the plasmid increases, given that the availability of mutants is dependent on the population size.

Note that the results of this study correspond to a limited number of possible parametric landscapes; however, our intention was to use the parameters that frequently influence plasmid and bacterial dissemination in hospital settings. In any case, a major advantage of the membrane computing modeling technology is its *scalability*, allowing to include many different parametric values simultaneously in the model at the various hierarchical levels considered in particular ecosystems in workload and scope. How changes in a “piece” (the plasmid) contributes to create a particular “pattern” in a nested system of biological units, expanding from the genes and cells to the communities of human hosts and environments, is certainly one of the challenges of modern research on antibiotic resistance (6, 58, 59).

## Materials and Methods

### Computing model

All computational simulations were performed using an updated version of the Antibiotic Resistance Evolution Simulator (ARES), which is a P system software implementation for modeling antibiotic resistance evolution (11, 13). The current version of ARES (2.0) can be freely downloaded at https://sourceforge.net/projects/ares-simulator/. The original ARES website http://gydb.uv.es/ares, offers information on the rules and parameters currently used by ARES and facilitates customer generation of specific scenarios to model the evolution of antibiotic resistance.

### The basic model application: quantitative structure

A detailed account of the main features of our model’s quantitative structure is available in our previous publication (12), a summary of which is presented below.

### Hospitalized hosts, admissions, and discharge rates in the population

The number of hosts in the hospital reflects an optimal proportion of 10 hospital beds per 1000 individuals in the community (https://data.oecd.org/healtheqt/hospital-beds.htm). The hospital has 100 occupied beds and corresponds to a population of 10,000 individuals in the community. The admission and discharge rates from hospital are equivalent to 3–10 individuals/10,000 population/day (http://www.cdc.gov/nchs/data/nhds/1general/). In the basic model, 6 individuals from the community are admitted to the hospital and 6 are discharged from the hospital to the community per day (approximately at 4 hour-intervals). Approximately 75% of the patients stay in the hospital between 6 and 9 days.

### Transfer of bacterial organisms between hospitalized hosts

We used a “contagion index” of 5% (for every 100 hospitalized patients, 5 “donors” transmit bacteria to another 5 “recipients” per hour. Bacterial transmission includes the spread of normal microbiota. The hospital is surrounded by a community of healthy individuals, occasionally admitted to the hospital, with a “contagion index” of 0.01%.

### Exposure to antibiotic agents

We considered 3 types of commonly used antibiotics to be employed during a 7-day treatment: aminopenicillins (AbA), third-generation cephalosporins, as cefotaxime (AbC), and fluoroquinolones (AbF). In the basic model, 20% of the individuals in the hospital compartment are under antibiotic exposure each day. Antibiotics AbA-AbC-AbF are employed in the hospital at a proportion (percentage) of 30-40-30, respectively. A single patient is treated with only one antibiotic, administered every 8 hours. After each dose is administered, all 3 (bactericidal) antibiotics induce after a decrease of 30% in the susceptible population after the first hour of dose exposure, and a 15% reduction in the second hour. In the community surrounding the hospital, 1.3% of individuals are undergoing antibiotic therapy; AbA-AbC-AbF is employed at a proportion of 75-5-20, respectively.

### The bacterial colonization space

of the populations of the considered clinical species (*E. coli, K. pneumoniae, Pseudomonas aeruginosa, Enterococcus faecium*) and other basic colonic microbiota populations is defined as the volume they occupy in the intestine. In natural conditions, the sum of these populations is estimated at 10^8^ cells per mL of colonic content. Clinical species constitute only 1% of the cells in each mL and have a basal colonization space of 1% of each mL of colonic content (or 0.01 mL). Other microbiota populations are considered a single ensemble. The colonic space occupied by these populations can change due to antibiotic exposure. AbA, AbC, and AbF reduce the intestinal microbiota 25%, 20%, and 10%, respectively. This space can be occupied by resistant populations of these human opportunistic pathogens; however, in the absence of antibiotic exposure, the colonic populations tend to return to the basal population size, which would occur in 2 months (60, 61).

### Populations’ operative packages and counts

To facilitate the execution of the model, we considered that 10^8^ cells in nature is equivalent to 10^6^ cells in the model. In other words, one “hecto-cell” (h-cell) in the model is an “operative package” of 100 cells in the real world. Given the high effective population sizes in bacteria, these 100 cells are considered a uniform population of a single cell type. For computational efficiency, we considered that each patient (in the hospital) or individual (in the community) is represented in the model by 1 mL of its colonized colonic space (approximately 3000 mL) and is referred to as a “host-mL”. Our results as therefore represented as “number of h-cells in all host-mLs” in most of the figures.

### Quantitative distribution of species and clones

In the basal scenario, the species distribution in these 1,000,000 cells (contained in 1 ml) was as follows: for *E. coli*, 860,000 cells, including 500,000 susceptible cells, 250,000 containing PL1-AbAR, 100,000 with the AbFR mutation, and 10,000 with the AbFR mutation and carrying PL1 (AbAR); for *E. faecium*, 99,500 cells susceptible to both AbA and AbF and 20,000 cells with chromosomal AbCR, AbFR, and CO1-AbAR (CO for *Enterococcus* conjugative element); for *K. pneumoniae*, 20,000 cells, with chromosomal AbAR and AbFR, also harboring PL3 (AbCR-AbAR); and *P. aeruginosa*, 500 cells containing PL3 (AbCR-AbAR) and chromosomal AbAR. At time 0, this distribution was identical in hospitalized and community patients.

### Bacterial multiplication rates

We considered the basal multiplication rate (corresponding to *E. coli* 0) to be equal to 1, in which each bacterial cell gives rise to 2 daughter cells every hour. Comparatively, the rates for *E. faecium, K. pneumoniae*, and *P. aeruginosa* were 0.85, 0.9, and 0.15, respectively. In these basic conditions, the acquisition of a plasmid or other incurs an cost of 0.06, while the acquisition of the AbFR mutation incurs a cost of 0.01 (1% reduction in growth rate) The number of cell replications will be limited by the available space (see above).

### Plasmids and antibiotic resistance types

For the sake of simplicity, this study considered 2 plasmids sharing the same partition system and therefore competing for replication when hosted in the same cell, with either 1) resistance to AbA, A for aminopenicillins) present in plasmid PL1, primarily hosted in *E. coli*, or (2) resistance to antibiotic C plus antibiotic A (AbC, C for third generation cephalosporins), primarily determined by the plasmid PL3, primarily hosted in *K. pneumoniae*. Note that some widely spread plasmids that encode extended-spectrum beta-lactamases (ESBLs) frequently carry genes for AbA resistance. In addition, mutational events lead to the emergence of chromosomal fluoroquinolone resistance (AbF, F for fluroquinolones). Organisms mutate to AbF at the same rate: 1 mutant for every 10^8^ bacterial cells per cell division. Note that *K. pneumoniae* has intrinsic (plasmid-independent) resistance to AbA. Although it was not analyzed in this study (but is included in the model), a clone of *E. faecium* resistant to AbA might transfer this resistance to a susceptible clone at a rate of 10^−4^ (62).

### Basal plasmid kinetic values

We established a basal set of plasmid kinetic values, based on reports in the literature (1,2, 19-23). In the simulation, modifications to each value were applied, maintaining the other values fixed; ultimately, more than one parameter was modified to ascertain the combined effects. The applied basal plasmid kinetic values were as follows: a) a plasmid transfer rate of 10^−6^ per hour; i.e., 1 in 100,000 or 1 million donor-recipient contacts that result in random and reciprocal cell-to-cell *E. coli*-*K. pneumoniae* transfer of the plasmid (10^−9^ from each of them to *P. aeruginosa*) every hour; b) a plasmid incompatibility rate of 2 plasmids/cell, indicating that, in the presence of a third plasmid, one of the three is stochastically removed; c) a rate of plasmid cost of 0.06; i.e., the bacterial growth rate is decreased by 6% when harboring plasmids); d) a rate of frequency of mutational plasmid cost compensation of 10^−5^; i.e., in 1 per 100,000 cells, a mutational event in the plasmid or bacterial genome decreases the plasmid cost by 50%; e) a rate of mutational events leading to significant fluoroquinolone resistance of 10^−8^; and f) a rate of plasmid segregation of 10^−5^, indicating that stochastically 1 cell among 100,000 eliminates the plasmid it contains.

## Acknowledgements

F. Baquero, M. Campos, and T. M. Coque were supported by EU Joint Programming Initiative JPIAMR2016-AC16/00043 (JPIonAMR-Third call on Transmission, ST131TS project), the Health Institute Carlos III of Spain (grants PI15-00818 and PI18-01942 and CIBER (CIBER in Epidemiology and Public Health, CIBERESP; CB06/02/0053)) and the Regional Government of Madrid (InGEMICS-C; S2017/BMD-3691); all of them cofinanced by the European Development Regional Fund [ERDF] “A Way to Achieve Europe”. A. San Millan was supported by the European Research Council under the European Union’s Horizon 2020 Research and Innovation Program (ERC grant agreement no.757440-PLASREVOLUTION).

## Supplemental Material

**Figure S1.**
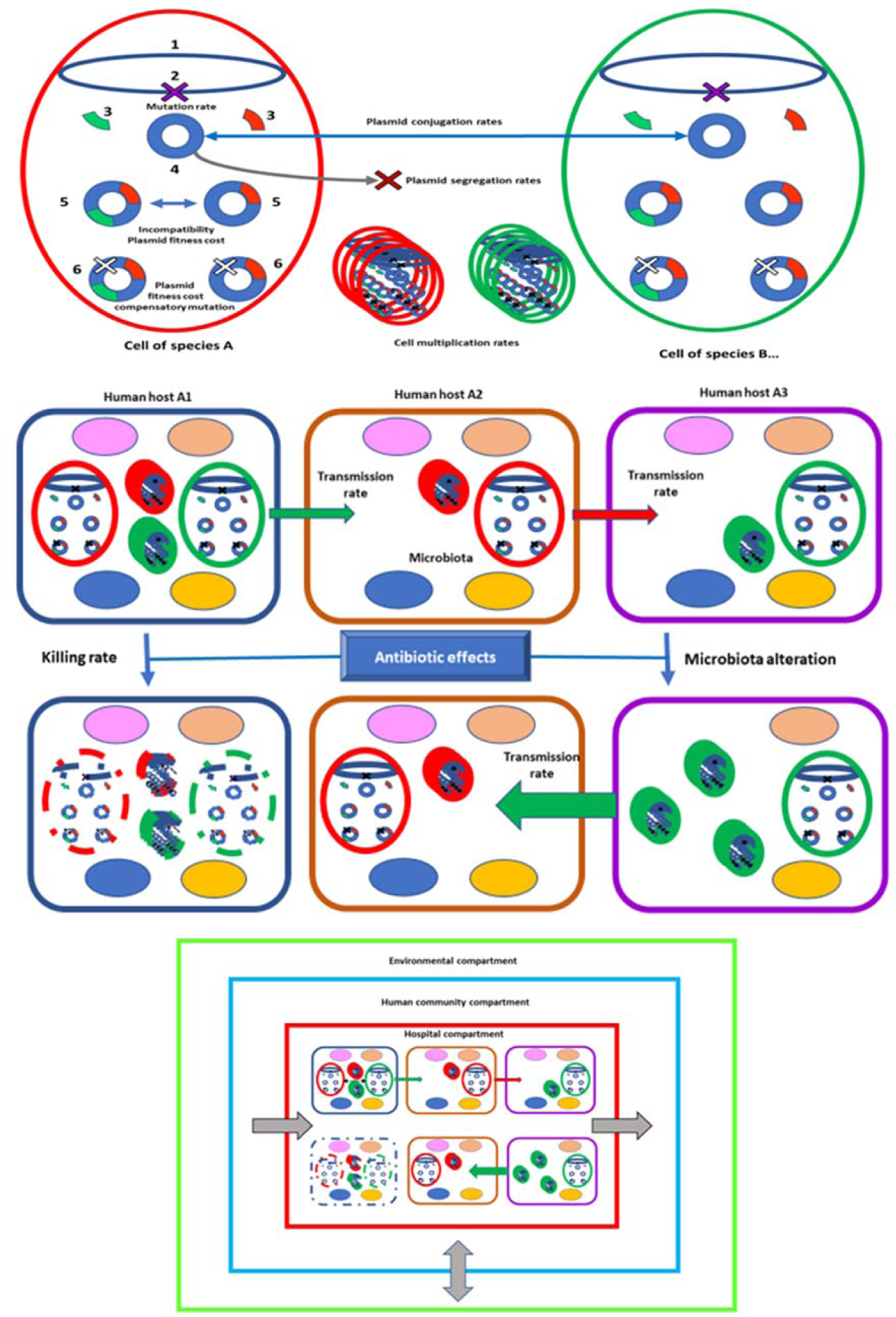
Simplified graphic schema of the membrane computation model. On the top panel, two bacterial cells of the same or different species (red and green circles) containing a chromosome (black oval, 1), where mutations might occur at variable rates (black X, 2); the absolute number of these cells might be modified. Inside each cell, different antibiotic resistance genes (red, green, 3), that might be harbored in plasmids of the same Inc groups (blue circle crown, 4). Plasmids might compete (incompatibility) inside the cell, eventually leading to the segregation of one of the replicons; also replicons might be submitted to random loss (X, 4), or impose a fitness cost to the host cell (5); however, cost-compensatory mutations can reduce this cost, restoring in part or totally the host-cell replication rate (6). Cells can replicate at different rates (cylinders of red or green ovals). Plasmids can be transferred between cells at different conjugation rates (horizontal black line). On the middle panel, each one of the squares with curved angles correspond to a different human hosts (different colors) where these bacterial cells are established; the colored ovals inside each host correspond (in a simplified way) to the different species in the microbiota. Bacteria harboring plasmids with resistance genes can be transferred from human to human hosts at variable rates (for instance, influenced by hospital hygiene or cross infection). Submitted to antibiotic exposure, different antibiotics can kill (eliminate) bacterial cells at certain rates, but bacteria might survive is they have resistance genes; note that other bacteria of the microbiota can also be eliminated by antibiotics, eventually increasing the population size of the surviving resistant bacteria (as the green cylinders down right), which can be transferred to new hosts. In the lower panel it is depicted that all these processes can occur inside a hospital (red square), or in the human community where the hospital is located (blue square); these compartments are linked by variable admission and discharge rates (horizontal grey arrows) that can also be introduced in the model; finally, the bacterial composition inside the human community is influenced by the interaction with the environment (green square). This figure reveals the multi-nested structure of units involved in antibiotic resistance.

